# Biases in volumetric versus surface analyses in population receptive field mapping

**DOI:** 10.1101/2024.09.02.610800

**Authors:** David Linhardt, Michael Woletz, Pedro M. Paz-Alonso, Christian Windischberger, Garikoitz Lerma-Usabiaga

## Abstract

Population receptive field (pRF) mapping is a quantitative fMRI analysis method that links visual field positions with specific locations in the visual cortex. A common preprocessing step in pRF analyses involves projecting volumetric fMRI data onto the cortical surface, often leading to upsampling of the data. This process may introduce biases in the resulting pRF parameters. To investigate this, we present CON-pRF, a fully containerized pipeline for pRF mapping analysis that is designed to maximize reproducibility in pRF mapping studies. Using this pipeline, we compared pRF maps generated from original volumetric with those from upsampled surface data. Our results show substantial increases in pRF coverage in the central visual field of upsampled data sets. These effects were consistent across early visual cortex areas V1-3. Further analysis indicates that this bias is primarily driven by the non-linear relationship between cortical distance and visual field eccentricity, known as cortical magnification. Our results demonstrate that reproducible analysis pipelines enable the detection of potential biases introduced by varying processing steps, particularly when comparing across differently processed datasets.

**Key Points:**

- Spatial upsampling increases pRF coverage in the fovea due to enhanced CNR and cortical magnification.
- The study highlights the need for careful consideration of data processing steps.
- CON-pRF is a containerized pipeline for enhanced reproducibility in pRF analysis.

## Introduction

The retinotopic organization of the visual system is characterized by the mapping of adjacent positions in the visual field to adjacent positions within the visual cortex (Wandell, Dumoulin and Brewer, 2007). Functional MRI (fMRI) enables us to assess only the collective neural activity within parcellated voxels of the human cortex rather than responses of individual neurons (Logothetis, 2008). However, population receptive field (pRF) mapping enables estimating the combined receptive fields of a population of neurons within a voxel (Dumoulin and Wandell, 2008). This approach has been used to assess the organization of the human visual system, showing high reproducibility across repeated acquisitions both in healthy subjects (Van Dijk *et al*., 2016) and in data from subjects with scotoma (Linhardt *et al*., 2022).

In the last few years, standardization of MRI data analysis has been a major topic in the brain mapping community and has recently seen considerable advancements being made. One of the major steps in this endeavor was the introduction of the Brain Imaging Data Structure (BIDS) format (Gorgolewski *et al*., 2016) which paved the way for standardized pipelines. Tools like fMRIPrep (fmriprep.org; Esteban *et al*., 2019) have been pivotal in providing containerized solutions for the processing of fMRI data, ensuring consistency across varying computational environments. Other containerized software packages aim for easy to use and reproducible software solutions, for example in the field of diffusion MRI and tractography (Lerma-Usabiaga *et al*., 2023). On a broader note, software such as Neurodesk allows running most available neuroimaging analysis software in a controlled software environment to ensure reproducibility (Renton *et al*., 2024).

In the field of pRF, the tool *prfanalyze* (Lerma-Usabiaga *et al*., 2020) has similarly streamlined and standardized the workflow by encapsulating pRF analysis steps in a container, allowing for data analyses with various tools other than prfanalyze, including vistasoft, AFNI, and popeye. Together with *prfsynth*, a software container capable of synthesizing artificial pRF mapping time courses, this ensures reproducible validation of different pRF analytical tools. Significantly, this package introduced standardized pRF input and output formats, allowing for direct comparison of results derived via a range of analysis suites (Lerma-Usabiaga *et al*., 2020). In addition to standardizing pRF analyses, *prfanalyze* also simplifies the development of novel tools for processing and analyzing pRF data, as well as facilitating their comparison with other available methods.

Such comparisons across methods are essential to assess the effects of study-specific pipelines on pRF results in order to reveal potential biases caused by the respective preprocessing and analysis variants, the parameter selection and the software environments in which these analyses are run. In order to be able to systematically examine these issues, we have developed CON-pRF, a reproducible containerized pipeline aiming to standardize the pRF analysis from data acquisition to result visualization. It is built by combining published and newly developed containers with the principle of separating every step of the preprocessing, analysis and visualization pipeline into a separate container. *CON-pRF* is an extension of the *prfanalyze* package and closes the gap between preprocessing and analysis with capabilities for the masking of visual areas and visualizing of model fitting results. The CON-pRF approach allows for comprehensive control of all stages of the preprocessing and analysis steps and extends the reproducibility beyond the initial phase of data acquisition and analysis. Herein, we use this novel package to assess effects of surface-based analysis on pRF results, an important topic in current pRF mapping.

As in other fields of fMRI analysis, there is a growing trend in pRF mapping studies to transition from a traditional volumetric voxel space (Dumoulin and Wandell, 2008; Pawloff *et al*., 2019; Linhardt *et al*., 2021; Prabhakaran *et al*., 2021) to a projection of the volumetric data onto the gray/white matter surface vertex space (Infanti and Schwarzkopf, 2020; Morgan and Schwarzkopf, 2020; Farahbakhsh *et al*., 2022; Urale *et al*., 2022; Himmelberg *et al*., 2023). Also, publicly available pRF mapping datasets such as the NYU dataset (Himmelberg *et al*., 2021) or the human connectome project (HCP) retinotopy data (Benson *et al*., 2018) published their results projected on a *freesurfer* surface template. While the volumetric approach allows for analysis with minimal preprocessing steps, surface analysis facilitates visually appealing presentation of results on the cortical surface as well as the possibility of unifying vertices across a large group of subjects e.g. for machine learning applications (Ribeiro, Bollmann and Puckett, 2021). Despite the benefits of surface analysis, there are also potential disadvantages, such as the interpolation of volumetric time-series data, which can introduce additional smoothness and other biases to the results. In this study, we aim to closely examine these potential biases, emphasizing the need for caution when comparing outcomes from volumetric and surface analyses.

We found a strong bias towards higher coverages in the central visual field compared to the periphery for the early visual areas V1-3. We hypothesize that this bias might be due to a combination of three causes: (1) increased contrast-to-noise (CNR) in surface compared to volumetric data due to averaging and smoothing; (2) spatial upsampling (more vertices than original voxels) during cortical projection yielding a greater number of pRF centers; and (3) spatial upsampling effects on the results due to the nonlinear mapping from cortex to visual field space (cortical magnification; Daniel and Whitteridge, 1961).

To further understand the underlying reasons for this bias, we ran a set of analyses. To check for hypothesis (1), we eliminated the effect of CNR increase by repeating the original analysis on simulated data with varying noise levels. To check for hypothesis (2), we ran two analyses: first, we compared surface and volumetric data with the same number of data points by random sampling of the surface data set; second, we compared the original surface to a subsampled surface data set. To test for hypothesis (3), we reproduced the bias by directly upsampling the original 2mm isotropic voxels to 1mm isotropic voxels.

## Methods

### Participants

A total of 30 right-handed volunteers (23.8±3.0 years; 16 female) were measured in two scanning sessions, 8.0±1.5 days (mean ± standard deviation) apart. Participants had normal or corrected-to-normal vision and no known ocular pathologies, gave written informed consent and were financially reimbursed. The study protocol was approved by the ethics committee of the BCBL and complied with the guidelines of the Helsinki Declaration.

### Data acquisition

All scans were performed on a SIEMENS Trio 3T scanner using a 32-channel head coil. To minimize movement throughout the scanning sessions, participants’ head motions were restricted within the coil using extensive cushioning. Participants were able to see the rear-projection screen located outside the scanner bore through a mirror, mounted on top of the head. Functional full-brain data were acquired using the CMRR multiband EPI sequence (Moeller et al., 2010) with the following parameters: voxel size=2x2x2Lmm; TR/TE=1883/30.8 ms; 72 slices; flip angle=75°; matrix size=96x96; FoV=192x192Lmm; phase enc=anterior-posterior; multiband=3; grappa=1. Slices of the full-brain images were aligned parallel with the corpus callosum. Within each of the two scanning sessions, four runs were acquired per subject, each comprising 135 volumes and lasting 4:14 minutes. In addition, full-brain anatomical T1-weighted scans were acquired with a multiecho (ME) MPRAGE sequence with TE-s = 1.64, 3.5, 5.36, and 7.22 ms, TR = 2.530 ms, flip angle = 7°, field of view (FoV) = 256 × 256 mm, 176 slices, and voxel size = 1 mm isotropic.

During the functional scans, subjects were presented with a bar aperture (bar width=1.95°) moving through the visual field in eight different directions revealing an 8Hz reversing checkerboard pattern. The stimulated field of view was circular, with a radius of 7.8°. Subjects were instructed to maintain central fixation. Attention was assessed by requiring subjects to report the number of color changes of the central fixation disc overlaid on the bar stimulus. Further information about participants and data acquisition can be also found in (Lerma-Usabiaga, Carreiras and Paz-Alonso, 2018)

### CON-pRF preprocessing and analysis pipeline

The fMRI data was prepared and analyzed using CON-pRF, a reproducible pipeline based on the concatenation of containerized analyses (see Figure 1 and Table 1). Example subject data processed with the presented CON-pRF analysis pipeline can be found at osf.io/seh6b. It includes all necessary data and scripts as well as the respective analysis results.

**Figure 1.**
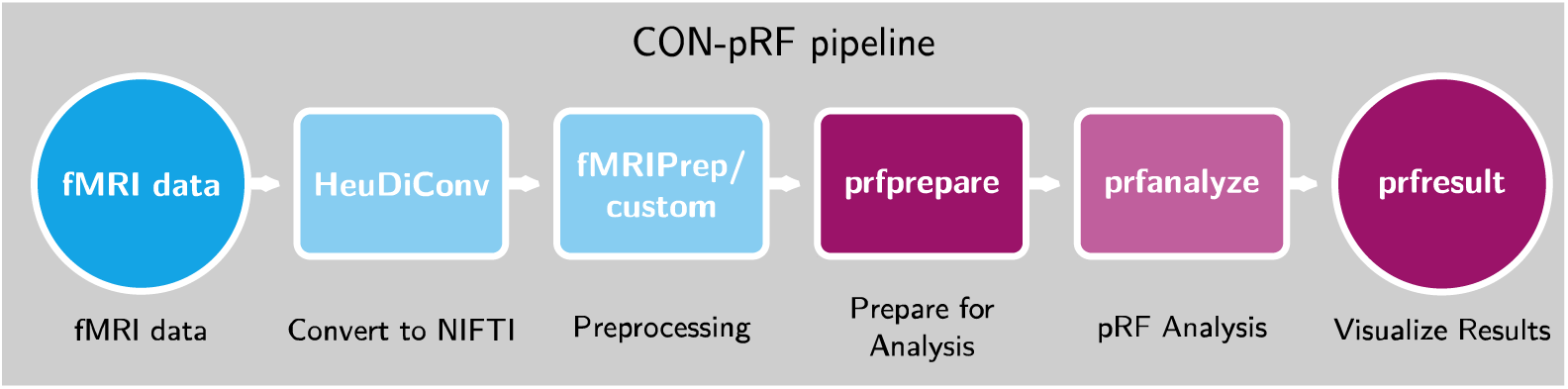
Flow-chart of the CON-pRF containerized pipeline for pRF analyses. The CON-pRF pipeline includes all the MRI preprocessing, pRF analysis and result visualization stages, starting with the original DICOM scanner data. **HeudiConv** converts DICOMs to NIfTI and curates them into BIDS format. **fMRIPrep** minimally preprocesses the data. **Prfprepare** takes the preprocessed functional data in anatomical or surface space and prepares it for the pRF analysis: it selects only the voxels/vertices the user wants to analyze, and prepares the stimuli. **prfanalyze** performs pRF analysis using different algorithms (such as vistasoft, AFNI, or prfanalyze) and was only slightly modified for a seamless experience. **Prfresult** plots the results as coverage maps or surface overlays. Blue color indicates pre-existing parts of the pipeline, while purple color indicates novel or modified (pale) containers.

**Table 1:** Overview of all containers comprising the CON-pRF pipeline, including their corresponding GitHub repository links.

|  |  |
| --- | --- |
| HeuDiConv | https://github.com/nipy/heudiconv |
| fMRIPrep | https://github.com/nipreps/fmriprep |
| prfprepare | https://github.com/fmriat/prfprepare |
| prfanalyze | https://github.com/vistalab/prfmodel |
| prfresult | https://github.com/fmriat/prfresult |

Acquired data was converted to NIfTI format using the containerized version of HEUDICONV v0.11.3 (github.com/nipy/heudiconv). The resulting NIfTi files used in this manuscript were obtained with preprocessing performed using fMRIPrep 23.0.1 (Esteban *et al*., 2019) and prfprepare (code: github.com/fmriat/prfprepare; container: hub.docker.com/r/davidlinhardt/prfprepare).

### Anatomical data preprocessing

T1-weighted (T1w) images were corrected for intensity non-uniformity (INU) with N4BiasFieldCorrection (Tustison *et al*., 2010), distributed with ANTs 2.3.3 (Avants *et al*., 2008). An anatomical reference map was computed after coregistration of two INU-corrected T1w images (one per session) using mri_robust_template (FreeSurfer 7.3.2, (Reuter, Rosas and Fischl, 2010)). This T1w-reference image was then skull-stripped with a Nipype 1.8.5 (Gorgolewski *et al*., 2011) implementation of the antsBrainExtraction.sh workflow (from ANTs), using OASIS30ANTs as target template. This image was used for brain tissue segmentation into cerebrospinal fluid (CSF), white-matter (WM) and gray-matter (GM) using FAST (FSL 6.0.5.1:57b01774, (Zhang, Brady and Smith, 2001)). Brain surfaces were reconstructed using recon-all (FreeSurfer 7.3.2, (Dale, Fischl and Sereno, 1999)), and the brain masks estimated previously were refined with a custom variation of the method to reconcile ANTs-derived and FreeSurfer-derived segmentations of the cortical gray-matter of Mindboggle (Klein *et al*., 2017).

### Functional data preprocessing

For each of the eight BOLD runs per subject (two sessions with four runs each), the following preprocessing was performed. First, a reference volume and its skull-stripped version were generated by aligning and averaging one single-band reference image (SBRef). Head-motion parameters (transformation matrices) with respect to the BOLD reference were estimated before any spatiotemporal filtering using MCFLIRT(FSL 6.0.5.1:57b01774, (Jenkinson *et al*., 2002)). BOLD runs were slice-time corrected to 0.906s (center of slice acquisition range 0s-1.81s) using the 3dTshift routine included in AFNI (Cox and Hyde, 1997). BOLD time-series were resampled by applying the transforms to correct for head-motion. These resampled BOLD time-series will be referred to as preprocessed BOLD in original space, or just preprocessed BOLD.

BOLD reference data was then co-registered to the T1w reference image using bbregister (FreeSurfer) which implements boundary-based registration (Greve and Fischl, 2009). Co-registration was configured with six degrees of freedom. First, a reference volume and its skull-stripped version were generated using a custom methodology of fMRIPrep. The BOLD time-series were resampled onto the fsnative and fsaverage surfaces, where fsnative is the individual subject surface representation and fsaverage is based on the MNI305 surface representation. All resamplings were performed with a single interpolation step by composing all the pertinent transformations (i.e. head-motion transform matrices, susceptibility distortion correction when available, and co-registrations to anatomical and output spaces). Gridded (volumetric) resamplings were performed using antsApplyTransforms (ANTs), configured with Lanczos interpolation to minimize the smoothing effects of other kernels (Lanczos, 1964). Non-gridded (surface) resamplings were performed using mri_vol2surf (FreeSurfer). Many internal operations of fMRIPrep use Nilearn 0.9.1 (Abraham *et al*., 2014), mostly within the functional processing workflow.

### prfprepare

To facilitate the analysis of minimally preprocessed data from fMRIPrep or other custom preprocessing pipelines with pRF analysis, we have developed a novel container (github.com/fmriat/prfprepare). This container bridges the gap between preprocessed data and the containerized solution that offers various pRF mapping tools, initially presented in (Lerma-Usabiaga et al., 2020). In prfprepare, the preparation of data for subsequent analysis involves three key steps.

First, log files and stimulus images from the vistadisp stimulus presentation suite (github.com/vistalab/vistadisp) are used to generate a stimulus representation in NIfTI format. Alternatively, the stimulus file can be provided if a different stimulus presentation suite was used. The container automatically links the correct tasks to the corresponding stimulus files.

Second, prfprepare container prepares the fMRI data for analysis, either in volume space (original voxel) or in surface space (Freesurfer’s *fsnative* space), as provided by fMRIPrep. It is also possible to restrict the analysis to visual regions only. Masking can be done at the voxel/vertex level and using two different atlases: Wang (Wang et al., 2015) or Benson (Benson et al., 2014). For this masking, the Neuropythy tool (Benson and Winawer, 2018) (github.com/noahbenson/neuropythy) is used. The masked data is then saved as a single NIfTI file per subject, run and hemisphere, containing all time courses of the masked voxels/vertices to be used for pRF analysis. Additionally, a sidecar JSON file is saved for each defined region of interest (ROI), specifying the indices for the corresponding ROI and their positions in the original volumetric or surface file. This information is later used to reconstruct the original file structure and to obtain data grouped by ROI.

Lastly, the prfprepare container organizes all output files following the BIDS guidelines for file naming in a designated folder structure. This ensures a seamless experience when executing subsequent analyses. Whenever parameters are modified, a new analysis folder is automatically created to prevent overwriting of old data.

### prfanalyze

prfanalyze enables the user to analyze real or synthetic data using different pRF analysis tools. It was part of a set of containers originally published in (Lerma-Usabiaga et al., 2020) (github.com/vistalab/PRFmodel), in which the user could synthesize ground-truth retinotopic data (prfsynthesize), analyze it with a set of existing tools (prfanalyze) and visualize the results (prfresults). For CON-pRF, the prfanalyze container has been updated to facilitate the interaction with the new prfprepare and the updated prfresult. The implemented algorithms are the same, namely:

● Vistasoft (github.com/vistalab/Vistasoft)
● analyzePRF (github.com/cvnlab/analyzePRF)
● popeye (github.com/kdesimone/popeye)
● AFNI (github.com/afni/afni).

All analysis containers share a common base image, using the same input and output data schemes. This standardized base image simplifies the implementation of new analysis tools. Both the base image and the analysis tool containers have been adapted to seamlessly analyze subject data processed with prfprepare, creating an integrated analysis pipeline. Users can easily switch or compare different tools as unified scripts for processing and the analysis of the data across all analysis tools are used.

### prfresult

The prfresult container is a from-scratch reimplementation of prfreport. This new version first reconstructs the results in the original functional space, regenerating the whole brain (surface or volumetric depending on how the data was analyzed). Second, it generates visual field coverage plots and cortex overlays for the obtained population receptive field (pRF) parameters. The overlays can be created in two formats: a rotating inflated surface animation as a gif file and an image of a sphere encompassing the entire occipital cortex. The container is based on a python class (github.com/dlinhardt/prfclass) implemented as a container (github.com/fmriat/prfresult).

### Data analysis

As the direct mapping between the volumetric and surface analysis spaces cannot be established easily, we employed a comparison of the resulting visual field coverage maps. Visual field coverage maps (Amano, Wandell and Dumoulin, 2009) were created independently for every run and condition (volume and surface results) as follows. Every voxel/vertex is represented by a two-dimensional Gaussian function in visual field space. This Gaussian is defined using the fitted parameters for center position (x, y) and pRF size (σ) with a maximum height of one. The coverage map is calculated as the average value at every visual field location, taking into account all above threshold voxels. In Figure 1 coverage is represented by the color-coded value in the background, while gray dots show every pRF center fitted. If not stated otherwise, the data were thresholded using a minimum variance explained of 20%. Disparities between these coverage maps were calculated by evaluating Cohen’s d at each position in the coverage plot. For a robust group-level analysis, we employed a bootstrapping method across single-subject Cohen’s d values, executing 50 iterations with replacement. The analysis scripts for these analyses are available at github.com/garikoitz/prf2d3d.

### Simulations

In addition to the experimental pRF data, we also wanted to examine effects of CNR and pRF sizes on the resulting maps. For this, we created 30 artificial subject data sets, with the same number of data points as acquired in the experimental data and simulated results for volumetric and surface processing. pRF center positions were drawn from a Gaussian distribution in the visual field, centered at the fovea. Single-subject coverage plots were calculated for both conditions and Cohen’s d effects were calculated, mimicking the analyses in the experimental data. To isolate the effect of CNR differences, we kept the pRF size constant while reducing CNR in the volumetric data. To isolate the effect of different pRF sizes, we kept CNR constant while changing pRF size in the surface set by 10%.

## Results

Traditionally, pRF analysis has been performed voxel by voxel in a volumetric environment (one time series per voxel). In some recent analysis pipelines, it is, however, common to project the data onto the white matter surface (expressed as a mesh with vertices) and perform the analyses in a bi-dimensional environment (one time series per vertex). Working on the surface has many computational and visualization advantages. But these come at the expense of losing layer information as information at the voxel level is collapsed onto the surface through averaging across different cortical depths. Depending on the size of the voxels and the chosen surface resolution, this transformation often entails upsampling the data (more vertices than original voxels). In the presented experiment, the given functional data has been measured with 2mm isotropic resolution. The conversion to the surface resulted, across subjects, in approximately 4.4 times more vertices on the surface of V1 than the number of voxels in the volumetric data. This ratio is increased even further when pRF parameters are fitted and thresholded by goodness-of-fit (variance explained). After applying the threshold to the two results independently, there were 5.8 times more surviving vertices in the surface V1 than voxels in the volumetric V1.

Figure 2 illustrates the initial bias we found when comparing volumetric and surface results. In Figure 2A we show the V1 coverage maps for both surface and volumetric results in one representative subject. It can be seen that the number of pRF centers (gray dots) present in the surface results is greatly increased compared to the volumetric variant. In Figure 2B, we compare surface-volumetric coverage for visual areas V1, V2, and V3 at the group level (N=30). The bootstrapped effect size of the difference (Cohen’s d) is shown for every location of the visual field (represented as a mesh of 128x128 pixels). In particular, the foveal part of the visual field yielded higher coverage for the results corresponding to the surface representation. More peripheral areas of the visual field show the opposite effect. Importantly, effect sizes increase in higher visual areas (V2, V3).

**Figure 2.**
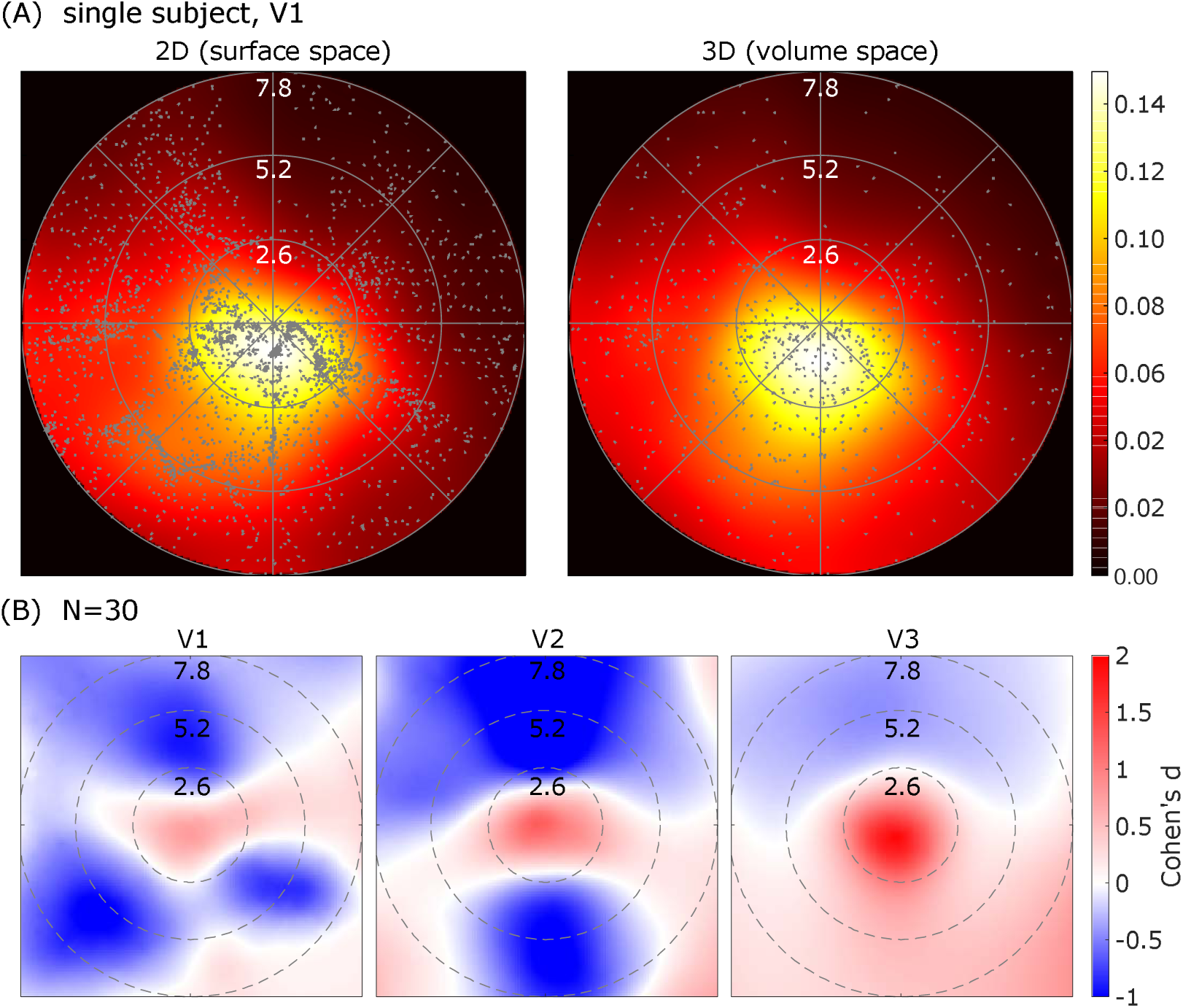
Coverage plots based on above-threshold voxels/vertices (10% variance explained threshold). **A.** Coverage plots for surface (left) and volumetric (right) data on V1 of a representative subject. There are substantially more pRF centers (gray dots) in the surface case, but the coverage looks similar, as expected when using mean values to calculate the coverage. **B.** In contrast, the group-based coverage surface-volume comparison (N = 30) shows that for the surface data there is a larger foveal coverage. Columns show the effect size (Cohen’s d) of the surface-volumetric coverage difference for V1, V2 and V3, with red meaning more coverage.

We derived three hypotheses for the effects observed: (1) increased contrast-to-noise (CNR) in surface compared to volumetric data due to averaging and smoothing; (2) spatial upsampling during cortical projection yielding a greater number of pRF centers; (3) spatial upsampling effects due to the nonlinear mapping from cortex into visual field space (cortical magnification).

To test for hypothesis (1), the effect of CNR differences in the two conditions was eliminated by using synthesized noiseless time series data (see Figure 3). We used previously calculated pRF parameters from the original analysis to compute the fitted model time series. We then substituted each voxel’s time series in the original volumetric data with these modeled noise-free time series. The procedure is similar to creating noiseless synthetic data where the spatial distribution of the ground truth parameters is known. Similar to the procedure used in the initial analyses, data was projected onto the surface using Freesurfer’s *mri_vol2surf* function. Results for the volumetric analyses should perfectly recover the same parameters used for the model creation (Lerma-Usabiaga et al., 2020).

**Figure 3.**
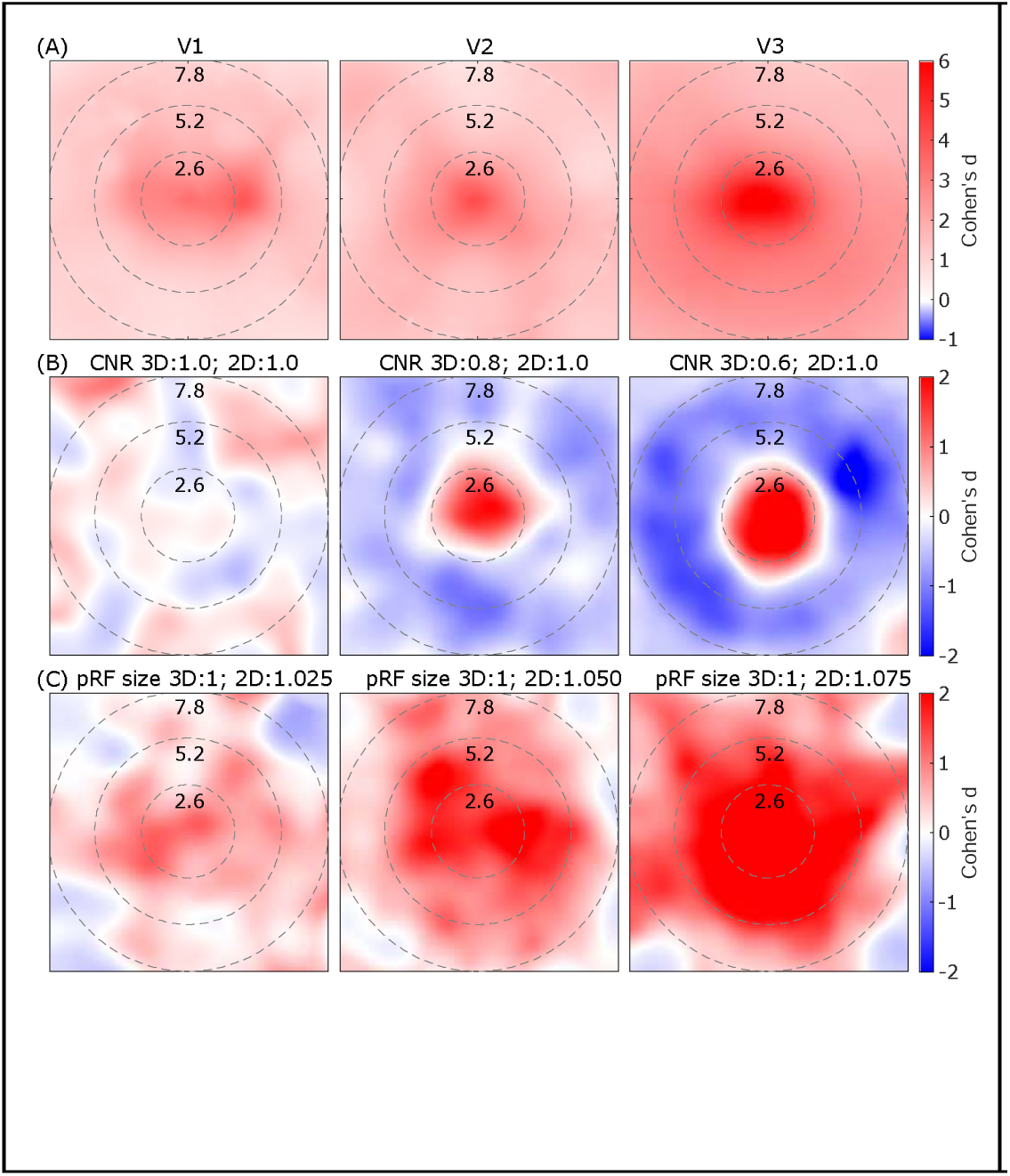
Group coverage surface-volumetric differences in noiseless data and simulations to check for CNR and pRF size effects. Bootstrapped (50 repetitions with replacement) effect size (Cohen’s d) representation of the surface-volume difference in field of view coverage depending on the different analyses. The comparisons were paired. A. Noiseless data in volumetric versus its projection to the surface. Only voxels with more than 50% variance explained were included. B. Simulations to examine CNR effects. pRF size between the two conditions was kept constant, while the CNR differences were increased left to right. With no CNR difference, the Cohen’s d map equals a random field (leftmost panel). With CNR differences, a clear bias towards higher coverage in the fovea and lower coverage in the periphery appears (center and right panels). C. Simulations to examine pRF size effects. CNR was equated in the two conditions, but the pRF size for the surface condition was increased left to right. With increasing difference a similar bias occurs, however the bias in foveal compared to peripheral areas was less pronounced.

The analysis was performed on all 30 simulated subject data sets. As data sets are noiseless, we used a threshold of 50% variance explained. Figure 3A shows the results of these analyses. It can be seen that coverage of surface analysis results is higher in the whole visual field throughout all early visual areas, as indicated by the absence of negative Cohen’s d values. However, the pattern of higher coverages in the central visual field seen in the experimental data is still clearly visible. As averaging and smoothing can lead to both a CNR increase and a pRF size increase, we ran two simulations to further isolate the two effects. Figure 3B shows the first simulation, maintaining a constant pRF size but different CNR values. We observe that there is no systematic bias in case of similar CNRs (left panel). When increasing the CNR differences, however, coverage increases in the fovea and decreases in the periphery (center and right panels). When maintaining a constant CNR but altering the pRF size (Figure 3C), we observe a similar bias as before (Figure 3A), but it is not as pronounced foveally and instead coverage increases across wide parts of the visual field.

In hypothesis (2) we conducted two different analyses to test whether the large difference in number of above-threshold voxels/vertices is the driving factor for the central coverage bias (see Figure 4). In the first analysis, the original volumetric results were compared with a same-sized random subsample of the surface data. Results from this comparison are shown in Figure 4A, revealing an almost identical effect as in the noiseless results (Figure 3A). In the second analysis, the original surface and the subsampled surface results were compared (Figure 4B), showing very similar results. These two analyses demonstrate that it is the underlying distribution of centers that drives the bias, and not the number of pRFs. In other words, the spatial upsampling operation itself is driving the bias.

**Figure 4.**
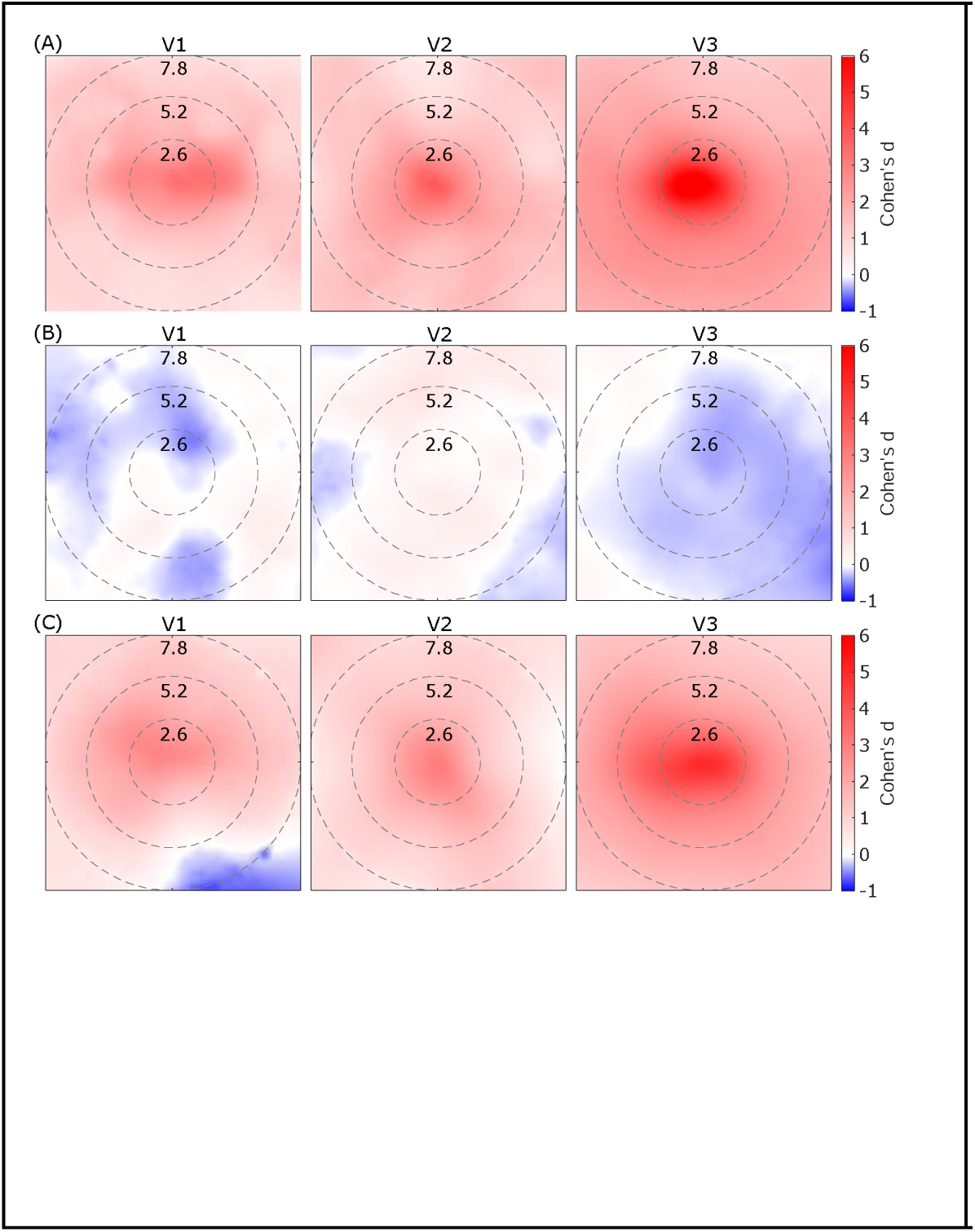
Upsampling effects on the differences between surface and volumetric processing. A. Subsampled surface results compared to the full volumetric results, with equal number of voxels/vertices. This analysis reproduces the central visual field bias. Thus, the number of data points is not the reason for the bias seen in the experimental results. B. Comparison of the surface data with a random subsample of the same data. No difference can be found in this analysis. **C.** Upsampled volumetric images (reslicing the voxels from 2 mm isotropic to 1 mm isotropic) compared to the original volumetric dataset. Here the same bias occurs as in panel A. To summarize, regardless of the number of data points, if there is upsampling (volumetric to surface or volumetric to volumetric), there is a foveal bias effect.

Hypothesis (3) states that the observed effect is due to the upsampling of data itself. For testing this hypothesis, the original volumetric data was spatially upsampled by reslicing from 2 mm to 1 mm isotropic resolution (see Figure 5). A trilinear interpolation kernel was used as implemented in the AFNIs 3dresample tool (Cox, 1996). Results for the upsampled data were compared to the low-resolution original volumetric data (Figure 4C). Again, a pronounced effect is observed, with the spatially upsampled data having greater coverage in the central visual field compared to the original not resliced data.

**Figure 5:**
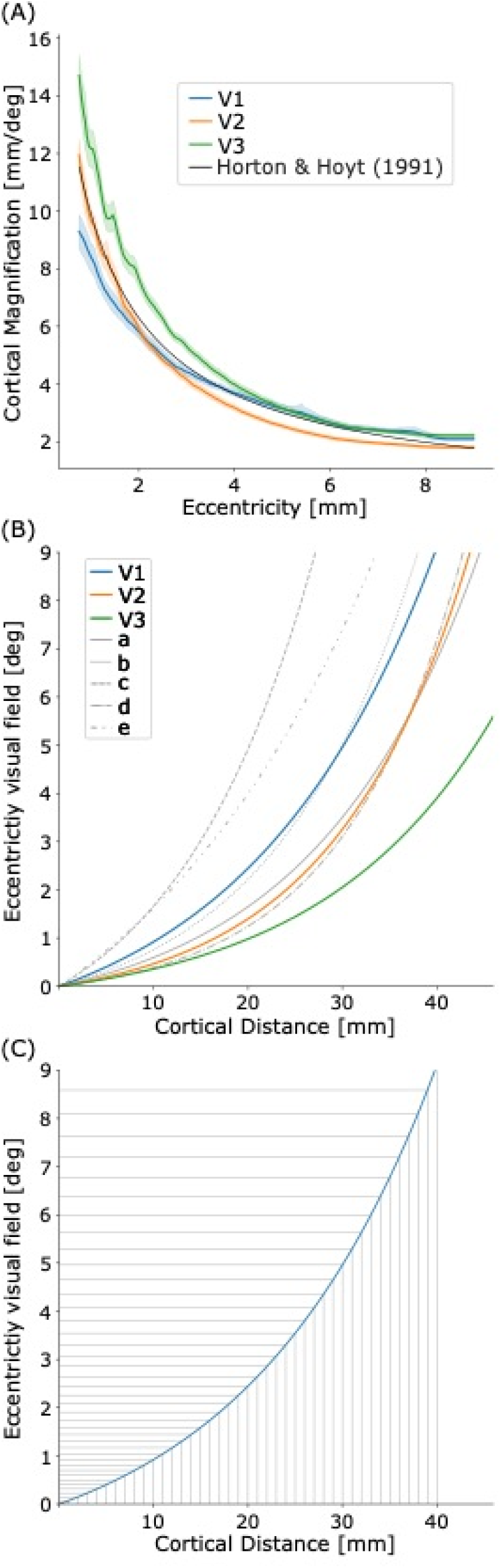
Cortical location function **A.** Linear cortical magnification function for the early visual cortex areas V1 (blue), V2 (orange) and V3 (green) with their 95% confidence intervals, overlaid with previously published results (black) from (Horton, 1991). **B.** Cortical location function on our experimental data pRF results for V1 (blue), V2 (orange), and V3 (green). For comparison, the gray lines represent V1 results as reported by Strasburger (2011) & (2022) and based on different studies (a: (Larsson and Heeger, 2006), b: (Duncan and Boynton, 2003), c: (Cowey and Rolls, 1974), d: (Schira et al., 2009), e: (Dougherty *et al*., 2003)). **C.** Cortical location function of the primary visual cortex V1 with an exemplary regular sampling of the cortical distance (same distance on x-axis). Due to the exponential relation with the position in the visual field, the sampling in the foveal areas is considerably denser than in more peripheral parts. When the data on the cortical level is linearly upsampled, this effect is intensified, leading to the effect observed in this study.

Based on our results, it seems clear that there is a combination of three effects driving the foveal bias, even though all of them originated in the data upsampling process. We isolated the smoothing and averaging processes in the first analysis and simulations (Figure 3), showing that biases are introduced by an increase in CNR and changes in pRF sizes.

Now we focus on the effect arising from non-linearities in the relationship between cortical distance and eccentricity in the visual field. This nonlinearity is characterized by the cortical magnification (Daniel and Whitteridge, 1961) of the visual system. It is modeled as an exponential function (Schwartz, 1977, 1980; Strasburger, 2022) and called the cortical location function (Equation 1)

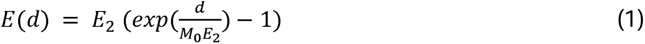

where *E* is the eccentricity in the visual field, and *d* is the distance from the occipital pole on the cortex. The constant M_0_ is called the cortical magnification factor and *E*_2_ the eccentricity value at which the inverse *M*^-1^ doubles the foveal value.

To estimate the cortical location function for our dataset, we first calculated the cortical magnification function shown in Figure 5A, based on the moving ring method (as described in github.com/noahbenson/cortical-magnification-tutorial). V1 data were thresholded at 10% variance explained and a maximum eccentricity of 9°. We fitted the linear cortical magnification function as shown in Equation 2 with the cortical scaling factor A (Dougherty *et al*., 2003; Schira *et al*., 2010; Harvey and Dumoulin, 2011).

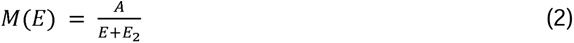

The resulting values for E_2_ [deg] are for V1: 1.4; V2: 0.5; V3: 0.5. We used this fit to calculate the cortical magnification factor M_0_ = M(0). The resulting M_0_ values [mm/deg] for the early visual cortex are for V1: 14.0; V2: 30.8; V3: 36.9. Plots of the cortical magnification factors over eccentricity are shown in Figure 5A with reference data from literature (Horton, 1991). Figure 5B shows the cortical location functions (Equation 1) for the obtained values for V1 (blue continuous line), V2 (orange continuous line), and V3 (green continuous line). For comparison, we plotted values taken from the literature in gray (Strasburger, Rentschler and Juttner, 2011; Strasburger, 2022). It can be seen that our experimentally obtained model for V1 is compatible with previously published results.

In Figure 5C, the cortical location function is shown in blue while gray lines visualize the described effect of stronger central visual field coverage. Linear sampling on the cortex (x-axis) leads to denser sampling in the central visual field compared to the periphery (y-axis) due to the exponential mapping.

## Discussion

In this work, we studied disparities in pRF-based visual field coverage maps when applying spatial transformation procedures: volume–to-surface and volume-to-volume upsampling. After the volume-to-surface transformation, we found an increase in pRF coverage in the fovea compared to the periphery (Figure 2). We characterized and isolated these effects in several analyses, and concluded that differences were due to a combination of two factors: increase in CNR (hypothesis 1) and cortical magnification (hypothesis 3). The number of data points, however, did not have an effect (hypothesis 2).

Upsampling data (both volume-to-surface and volume-to-volume) leads to increases in CNR due to averaging data from neighboring voxels. The averaging process smooths the time courses, also leading to an increase in pRF sizes fitted. The increase in CNR and pRF sizes partially explains the effect of increased foveal coverage observed in this study, through different mechanisms.

The first mechanism is related to the increase in CNR and its influence on the distribution of pRF centers in the visual field. The distribution of pRF centers is not uniform across the visual field, as there is a higher density in the foveal regions. When using fMRI and pRF mapping algorithms, noise introduces a broader probability distribution for each pRF center. Instead of being Gaussian, these probability distributions are skewed, pulling the centers toward more eccentric (peripheral) regions. As a result, the overall distribution of pRF centers across the visual field becomes wider, with more centers being inaccurately positioned toward the periphery. Due to the smoothing in the spatial upsampling operation, noise is reduced and the pRF center distribution is changed, again narrowing the distribution with a stronger peak and lower values in the periphery. When we now compare the unsmoothed with the smoothed distributions, we expect the results shown in Figure 2: more coverage in the center and less coverage in the periphery. Through simulations (Figure 3B) we isolated this partial effect.

A second mechanism is related to the fact that both types of upsampling are increasing the pRF size and therefore affecting the coverage. Temporal filtering using a low-pass filter, equivalent to smoothing of the time course, leads to peak widening and pRF size increases (Morgan and Schwarzkopf, 2020). This process systematically affects the whole visual field, but is more prominent in foveal regions. The effect of size increase was isolated in a separate simulation, maintaining constant CNR while varying pRF size (Figure 3C). This finding confirmed that pRF size plays a role in explaining the observed main effect, although its contribution is much smaller than the CNR increase.

In hypothesis (2), we considered the possibility that the difference in coverage is attributed to the substantial difference in voxel/vertex number between the upsampled and non-upsampled datasets. A priori, this should not be the case, given the way coverage maps are calculated, as values across all pRFs are averaged in every pixel of the visual field. To test this, we conducted two analyses. First, we randomly subsampled the surface data to match the size of the original volumetric data, showing results consistent with the noiseless comparison (see Figure 4A). Second, we compared the original surface data with the subset of the same surface data, and observed no discernible differences (Figure 4B). These findings suggest that variations in coverage are not related to the number of data points, but rather to the underlying distribution the samples are drawn from.

Our volume-to-volume upsampling analysis, and particularly the effects of cortical magnification (Daniel and Whitteridge, 1961), support our hypothesis (3) that both spatial upsampling and cortical magnification are contributing to the foveal coverage bias. However, the volume-to-volume upsampling analysis essentially replicates the original volume-to-surface effect without isolating the possible contribution of cortical magnification. To understand the underlying mechanism, we used the same experimental data to estimate the cortical location function (see Figure 5B), which links distinct positions on the cortex to positions on the visual field. We hypothesized that the experimental data would reflect this function. Our data showed a comparable magnification factor (*M*_0_, CMF) close to previously reported values with increasing cortical magnification factors when in higher visual areas. This increase in CMF explains our results showing a bigger effect for higher visual areas V2 and V3 than for V1 (see Figures 2, 3, 4). Previously reported CMF were not consistent when comparing visual areas, as they showed a reduction (Horton, 1991; Dougherty *et al*., 2003), an increase (Silva *et al*., 2018), and a difference that changes depending on the eccentricity (Schira *et al*., 2009, 2010). Experimental results obtained in the current study do not offer conclusive evidence in this regard, as cortical magnification functions for the early visual areas are crossing (Figure 5A).

Thus, the third contributing factor to the increased foveal coverage after both spatial upsampling methods, whether directly in the space of the volume or by projecting it onto the surface, is the exponential relationship between visual field eccentricity and cortical location (Keliris *et al*., 2019). This cortical location function and its effect is visualized in Figure 5C. When positions on the cortex (x-axis) are linearly upsampled, the effect is exponentially distributed in the visual field (y-axis). This leads to a non-linear increase in pRF center density in the central visual field relative to peripheral areas. Spatial upsampling of the data results in a more fine-grained sampling of the cortex and is equivalent to halving each interval on the x-axis of Figure 5C (gray lines). Though each subsampled step will have a similar distance to its neighbors on the x-axis, the exponential transformation will bring the corresponding position in the visual field slightly closer to the foveal neighbor than to the peripheral. Consequently, this causes an overall shift of the pRF center distribution towards the central visual field. Additionally, the exponential function increases the sampling density for the central visual field, while lowering it in the periphery. This effect describes the underlying distribution of pRF centers in the visual field, peaking in the center.

A limitation of this study is that the observed effect of increased foveal coverage was not demonstrated in data acquired with different voxels sizes. Future studies should address this question, although we consider this comparison complex. Together with changes in voxel size many other sequence parameters will change, making it challenging to isolate the voxel size effect alone.

Every choice made during data processing will influence final results, and researchers should be aware and consider tradeoffs associated with every step and how it may alter the final analysis results. While the processing steps used here might not substantially impact qualitative analyses such as GLM, they can meaningfully affect outcomes of quantitative methods such as pRF analyses. In modern fMRI analyses, there are several reasons for upsampling the data. In studies involving multimodal data, reslicing is used to unify resolution across modalities, i.e. bringing diffusion MRI (dMRI), quantitative MRI (qMRI) and functional MRI (fMRI) into the common T1w isotropic space (Lerma-Usabiaga, Carreiras and Paz-Alonso, 2018). Often, the other modalities are resliced to the high-resolution anatomical T1w template. Another common application of upsampling is the initially discussed situation where the data is upsampled when projecting volumetric time series to the surface mesh. Working on the surface has previously shown advantages (freesurfer.org; (Fischl, 2012)), and many current fMRI analysis pipelines include this step. In standard fMRI retinotopy analyses, the number of vertices in the surface approach is usually substantially higher from the original number of acquired voxels. Researchers must carefully interpret upsampled results, taking into account the systematic biases revealed in this study, as well as other potential biases introduced by data processing steps such as interpolation, smoothing, and the transformation of voxel data to surface space.

An additional contribution of this work is the presentation of a fully containerized analysis pipeline designed to streamline pRF mapping analysis with experimental data. This innovation builds upon previous efforts to containerize various pRF mapping tools (Lerma-Usabiaga *et al*., 2020), which emphasized the generalizability of tools based on reproducible and easy-to-use methods. Our new tools focus on providing a highly reproducible containerized analysis environment. The new main container is called *prfprepare*, and bridges the gap between minimal preprocessing, facilitated by fMRIPrep (Esteban et al., 2019), and advanced pRF analysis tools. Additionally, the *prfresult* container provides a platform for visualizing the results obtained from the analyses. The containers can be downloaded from DockerHub, and run using Docker or Apptainer/Singularity. The code and instructions for downloading and running the containers are available on github.com/fmriat/prfprepare and github.com/fmriat/prfresult.

## Conclusion

In conclusion, we have presented the new CON-pRF pipeline that is designed to ensure high reproducibility and traceability of the whole pRF mapping process. We have used CON-pRF to demonstrate that different processing strategies can have considerable impact on the results of pRF mapping analyses. As a quantitative fMRI technique, pRF mapping offers some advantages over classical GLM analyses. However, as our understanding of the technique evolves, we become aware of how different aspects of the analyses may influence the results. Our study revealed systematic biases introduced when transitioning from volumetric to surface analytical spaces in pRF mapping, or when spatially upsampling. These findings have significant conceptual and practical implications for the design, analytical decisions and interpretation of the results of future neuroimaging studies and warrant further investigation. The tools presented in this work will be valuable for future researchers investigating methodological or scientific questions in the field of pRF mapping.

## Acknowledgements

D.L., M.W. and C.W. were supported by the Austrian Science Fund (FWF; P35583) and D.L. by the EMBO Scientific Exchange Grant (Number 9415).

G. L.-U. was supported by the Spanish Ministerio de Ciencia e Innovación (IJC2020-042887-I; PID2021-123577NA-I00, RYC2022-035502-I; CEX2020-001010/AEI/10.13039/50110001103 3) and Basque Government (PIBA_2022_1_0014; KK-2023/00090).

P. M. P-A. was supported by grants from the Spanish Ministry of Science and Innovation (PID2021-123574NB-I00), from the Basque Government (PIBA-2021-1-0003), and from “la Caixa” Foundation (ID 100010434) under the agreement HR18-00178-DYSTHAL.

BCBL acknowledges funding from the Basque Government through the BERC 2022–2025 program and by the Spanish State Research Agency through BCBL Severo Ochoa excellence accreditation CEX2020-001010-S.

